# Study of *Staphylococcus aureus* Biofilm Ability to Destroy Bone Matrix in Absences of Host Immunity

**DOI:** 10.1101/2020.02.06.936799

**Authors:** ahmed Al Ghaithi, Sultan Al Mastari, John Husband, Mohammed al kindi, Atika Al Bimani

**Affiliations:** Ministry of Health Oman; Sultan Qaboos University Hospital; Sultan Qaboos University

## Abstract

**Purpose:** Osteomyelitis is an infectious bone process leading to bone necrosis and destruction. Published reports on pathogen biofilm thus far have focused on indirect bone resorption mediated by host cells and factors secondary to immune system activation. However, direct bone resorption due to biofilm pathogen has not been adequately studied yet. This study aims to investigate the effect of biofilm pathogen in ex-vivo human bones in the absence of the host immune response using Raman spectroscopy and Scanning electron microscopy.

**Methods:** Bone samples collected from patients who underwent knee replacement surgeries were inoculated with *Staphylococcus aureus* bacteria. Bacterial direct effects on the bone quality were then examined, at various time intervals, using Raman spectroscopy and scanning electron microscopy.

**Results:** Raman spectroscopy and scanning electron demonstrated the destruction of bone structure and drop in bone quality.

**Conclusion:** This experiment shows the direct effect of bacteria on bone during osteomyelitis in addition to the recognised destruction caused by the host immune system.

## Introduction

Osteomyelitis (OM) is bone inflammatory process secondary to an infectious organism leading to bone necrosis and destruction. Bone is a sterile organ resistant to bacterial colonisation. However, events like trauma, surgery, or hematogenous spread of virulent pathogen may disturb structural integrity and results in infection^(1)^. Several confounding factors may influence the inflammatory process of OM leading to diagnostic difficulties and treatment delays.

Due to the marked variability in osteomyelitis presentation and treatment, many researchers tried to investigate disease pathogenesis and introduce a controlled approach to active management. However, the primary focus of most of the published reports was a collection of infection-related parameters: haematological, radiological, histology or bacterial culture^(2)^. Some reports focused on the process of bone destruction due to immune system activation. However, there is limited emphasis on biofilm formation and pathogen-induced bone destruction^(3)^. Most causative pathogens can adhere to and grow on the surface of bone, joints and prosthesis causing bone resorptions. The pathophysiology of bacterium-induced bone resorption has not been identified. However, reports have suggested that pathogens such as *Staphylococcus aureus* express an array of cell surface and soluble molecules (e.g. acids and proteases) that have the potential to promote bone destruction at the infection site^(4)^. These molecules can either induce immune system activation or direct bone resorption due to their osteolytic properties.

Published reports on pathogen biofilm thus far have focused on indirect bone resorption mediated by host cells and factors secondary to immune system activation ^(3, 5, 6)^. However, direct bone resorption due to biofilm pathogen excretions has not been adequately studied yet. Therefore, we planned to investigate the effect of biofilm pathogen in ex-vivo human bones in the absence of the host immune response using Raman spectroscopy and Scanning Electron Microscopy. We chose Raman spectrometry because of its high ability to examine molecular changes of any material at ultrastructural levels. Recent reports have shown its effectiveness in evaluating bone quality in various pathologies^(7)^. It can provide information about mineral and organic content of bone at the molecular level.

## 2. Materials and Methods

### Bone samples preparation

Following informed written consent, bone cuts from patients undergoing total knee replacement were collected for the study. Sterile healthy cancellous bone samples measuring 1-2 cm^2^ were then cut using bone nibblers and stored under sterile conditions in Normal Saline at −20°C until the day of inoculation. Aseptic precautions were maintained throughout bone samples processing. Sterility of bone pieces was confirmed by culture swabs in blood agar medium for 48 hours just before the samples processing (none of the samples had any bacterial growth nor did any of the patients develop any infection subsequently).

A bacterial inoculum of *Staphylococcus aureus* was created from fresh overnight culture. Sterile normal saline was mixed with bacterial concentrate with a range of 1 x 10^6^ to 1 x 10^7^ colony forming units (CFU)/ ml. Bone pieces were then added to the mixture. The control bone samples were added to sterile normal saline. Each mixture contained 12 bone pieces. All bone samples were incubated at 37°C to allow bacterial growth and multiplication. Each week two bone samples from each mixture were taken for examination. Once examination was done, the samples were disposed of according to our hospital’s policy for human tissues.

### Bacterial growth and bone resorption examination

#### Scanning Electron Microscopy Setting

Bone samples were fixed in Karnovsky’s Electron Microscopy fixative (2.5% glutaraldehyde with pH 7.2) for 4 hours. Using washing buffer, they were twice washed each for 10 min. Samples were then post-fixed using 1% osmium tetroxide for 1 hour then dehydrated using serial concentrations of ethanol (25%, 75%, 95% and 99.9%). After that, samples were dried using Autosamdri-815^®^ (Rockville, MD, USA) mounted on Al stubs and then coated with gold using BioRad SEM coating system. Finally, samples were screened and micrographs were revealed using JEOL JSM-5600LV scanning electron microscope (JEOL Ltd, Tokyo, Japan).

#### Raman spectroscopy setting

Scanning of the bone samples was performed using a Raman spectrometer (I-Raman Ex spectrometer, BWTek, Newark, USA) with excitation laser diode operating at 1064nm. Spectra were recorded at (400-2000 cm^−1^).

Analysis of the obtained spectra was carried using BWSpec^®^ software (BWTek, Newark, USA). The software allowed automated spectra baselining and averaging to improve the signal to noise ratios. Manual identification of crucial Raman bands was carried out. The effect of the bacterial osteolytic process on the bone mineral content, the content of carbonate, collagen cross-linking, mineral and collagen fibril orientation were studied.

Statistical analysis of Raman parameters of inoculated bone samples compared to controls using Statistical Package for the Social Sciences 2016 (SPSS) (IBM Corporations, New York-USA) to determine the changes in the ultrastructure of examined bone architecture. Significant difference P value was set at < 0.05; Paired T test was used.

## Results

### Isolation of bacteria from analysed surfaces

Bacterial biofilms grown under the described conditions in Methods section were isolated from the surfaces of examined samples using swabs. Swabs were cultured in blood agar medium for 48 hours. Cultured swabs were positive for all inoculated samples for eight consecutive weeks compared to the negative culture for all control samples. The average bacterial count in petri-dish agar was 5 x 10^5^ CFU/ml.

### Raman spectrometry analysis

Using Raman spectrometry, we were able to study the micro-architectural changes in the bone. Standard bone spectra in figure 1A where phosphate represented at 958cm^−1^ peak and carbonate at 1070cm^−1^. Raman bands at 851, 873 and 917 cm^−1^ are characteristic of the collagen and hydroxyproline matrix and Raman band at 1001 cm^−1^ is distinctive of phenylalanine. Raman band located between 1210 and 1320 cm^−1^ relates to amide III. Figure 1B represents average Raman Spectra collected from both control and inoculated samples at eight weeks of incubation.

**Fig 1 A:**
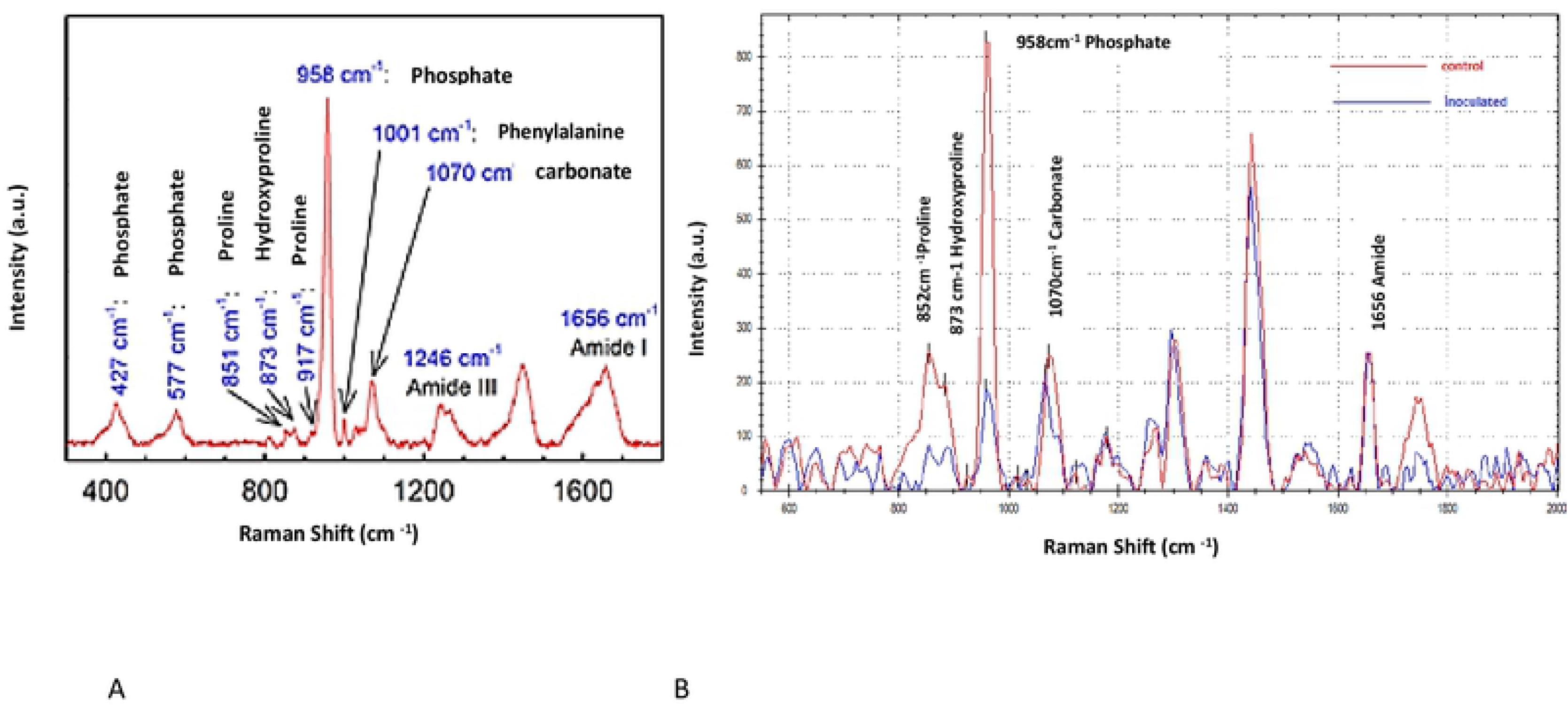
Typical Raman spectrum of healthy human bone with labelled waves’ peaks of interest. Fig1 B: average Raman spectra collected from both control and inoculated samples at eight weeks of incubation.

**Figure 1:**
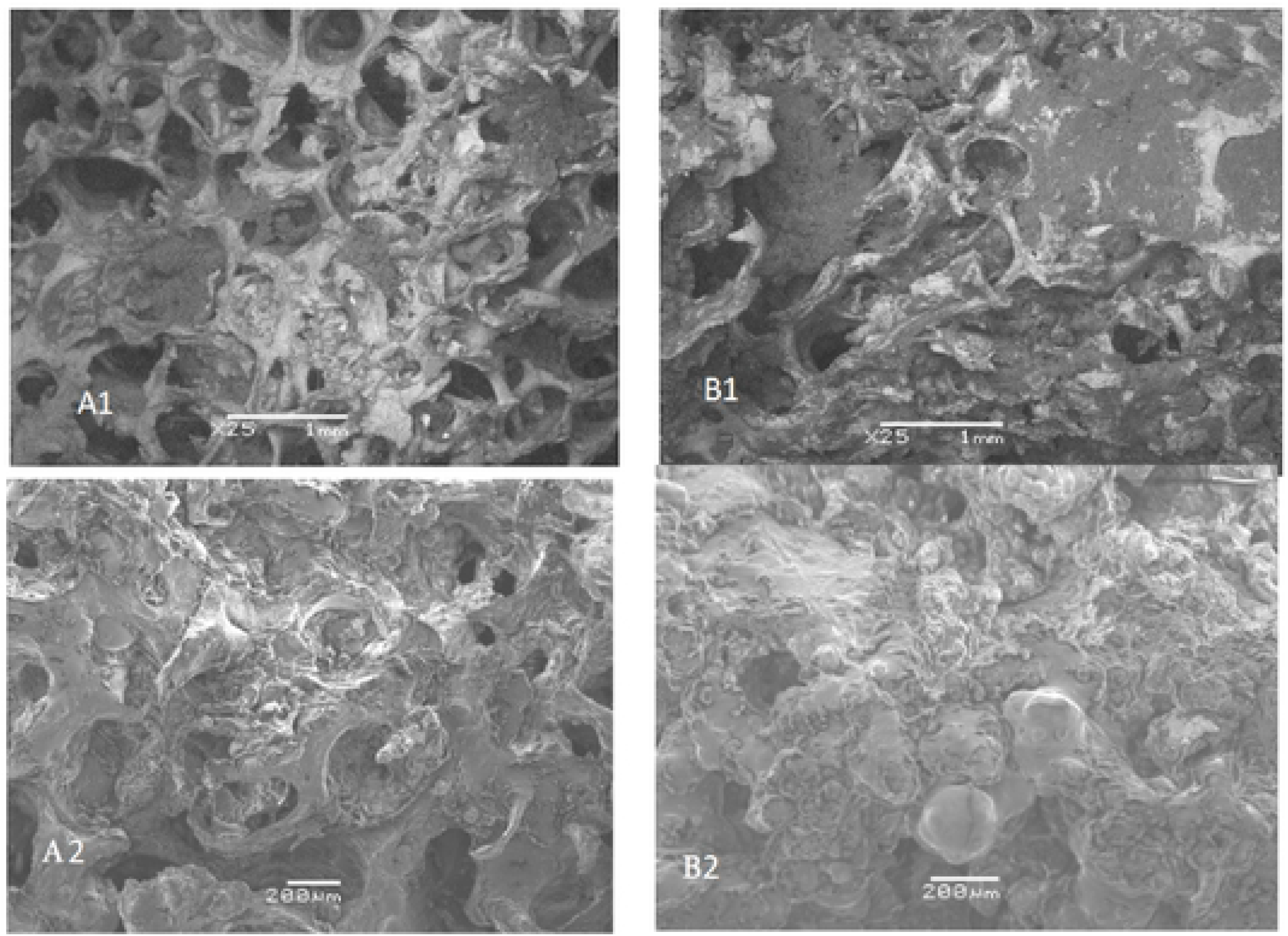
SEM images of control bone samples (fig. A1) and inoculated bone samples (fig. B1) at the end of week 8. Gross trabecular bone destruction by the *Staphylococcus aureus* biofilm is evident in B. Fig (A2 & B2) shows the samples trabecular architecture at higher magnification (200μm).

The carbonate to phosphate ratio represents the bone mineral content which was obviously increasing with time in the inoculated bone samples compared to the controls. This may indicate the change in bone crystallinity and transformation of a hard bone matrix to a brittle one. Control samples mean for Week 1 was 0.19 (SD 0.14; Min 0.17, Max 0.23), week two 0.23 (SD 0.073; Min 0.8, Max 0.28), Week three 0.22 (SD 0.48; Min 0.19, Max 0.26), week six 0.22 (SD 0.07; Min 0.19, Max 0.36), and Week eight 0.28 (SD 0.083; Min 0.19, Max 0.36). Mean ratio for inoculated samples Week one was 0.21 (SD 0.31; Min 0.16, Max: 0.23), week two 0.37 (SD 0.23; Min 0.14, Max 0.6), Week three 0.54 (SD 0.18; Min 0.31, Max 0.34), week six 1.18 (SD: 1.64; Min 0.17, Max 3.55), and week eight 1.51 (SD 1.26; Min 0.64, Max 2.38). The ratio increased by 619% in the inoculated samples compared to 47 % in the control group which was statically significant (P < 0.05).

Studying the mineral content of inoculated samples compared to control samples, a gradual drop of mineral content was recorded. The mineral to matrix ratio is represented by phosphate and carbonate bands relative to collagen band. The imbalance in this ratio indicates the destructive effect of bacterial biofilm contents on the bone trabecular surfaces. Therefore, this can have a negative impact on the bone mechanical properties resulting in fragility and fracture. These results are analogous to what happens in metabolic diseases of the bone where abnormal hormone levels can result in defective mineralisation and drop in the minerals to matrix ratio like osteoporosis and osteogenesis imperfecta^(7)^. Phosphate to amide mean ratio in control samples: week one 13.87 (SD 3.66; Min 11.28, Max 16.46), week two 12.41 (SD 1.05; Min 11.28, Max 13.37), week three 12.44 (SD 1.27; Min 11.28, Max 13.86), week six 9.95 (SD 1.45; Min 8.4, Max 11.28), week eight 6.25 (SD 0; Min 6.25, Max 6.25). Mean ratio for inoculated samples was: week one 13.20 (SD 3.49; Min 10.47, Max 15.67), week two 9.61 (SD 2.60; Min 5.03, Max 16.14) week three 5.45 (SD 0.42; Min 5.42, Max 6.15), week six 5.28 (SD 5.88; Min 1.13, Max 9.46) and week eight 2.39 (SD 0; Min 1.25, Max 6.25). 82% drop was observed in the inoculated samples compared to 55% in the control group but the difference was not statically significant (*P* > 0.05).

Carbonate to Amide I ratio, control samples mean was: week 1 was 0.04 (SD 0.0; Min 0.04, max 0.04), week two 0.0315 (SD 0.0105; min 0.02, max 0.04), week three 0.0299 (SD 1.45; Min 0.02, max 0.04), week six mean 0.023 (SD 0.00074; min 0.02, max 0.024) and week eight 0.0214 (SD 0.00692; min 0.01, max 0.03). Mean ratio for inoculated samples was: week one 0.06 (SD 0.05; min 0.03, max 0.1), week two 0.0196 (SD 0.007; min 0.01, max 0.02), week three 0.0129 (SD 0.005; min 0.01, max 0.02), week six 0.017 (SD 0.001; min 0.02, max 0.2), and week eight 0.0119 (SD 0.006; 0.01, max 0.02). 47% drop was observed in the inoculated samples compared to 80% in the control group but the difference was not statically significant (*P> 0.05*).

The organic content of examined bone samples was examined using the ratio of amide I Raman band at 1660 cm^−1^ to 1668 cm^−1^. The amide I band shift suggests rupture of collagen cross-linking which is a result of fibril scaffold deformation. This deformation can be attributed to the bacterial biofilm which affects the bonding within collagen fibril and binding of minerals to collagen. These results are similar to pathological bone resulting from hormonal changes or radiation^(8)^ where bone loses its mechanical properties and is fragile. Control samples mean ratio was: week one 3.89 (SD 5.4; Min 0.56, Max 19.23), week two 3.08 (SD 12.83; min −35.77, max 67.76), week three 1.6 (SD 0.46; min 1.33, max 2.16), week six 1.57 (SD 1.25; min 0.09, max 3.04), and week eight 1.09 (SD .75; min 0.57, max 1.62). Mean ratio for inoculated samples was: week one 3.149 (SD 1.32; min 1.62, max 3.91), week two 2.67 (SD 3.29; min 0.35 max 5), week three 1.34 (SD ?0.39; min 1.07, max 1.62), week six 1.31 (SD ?0.13; min 1.21, max 1.39), and week eight 0.56 (SD 0.01; min 0.054, max 0.65). 82% shift in amide band was observed in the inoculated samples compared to 72% in the control group but the difference was not statically significant (*P > 0.05*).

**Fig 2:**
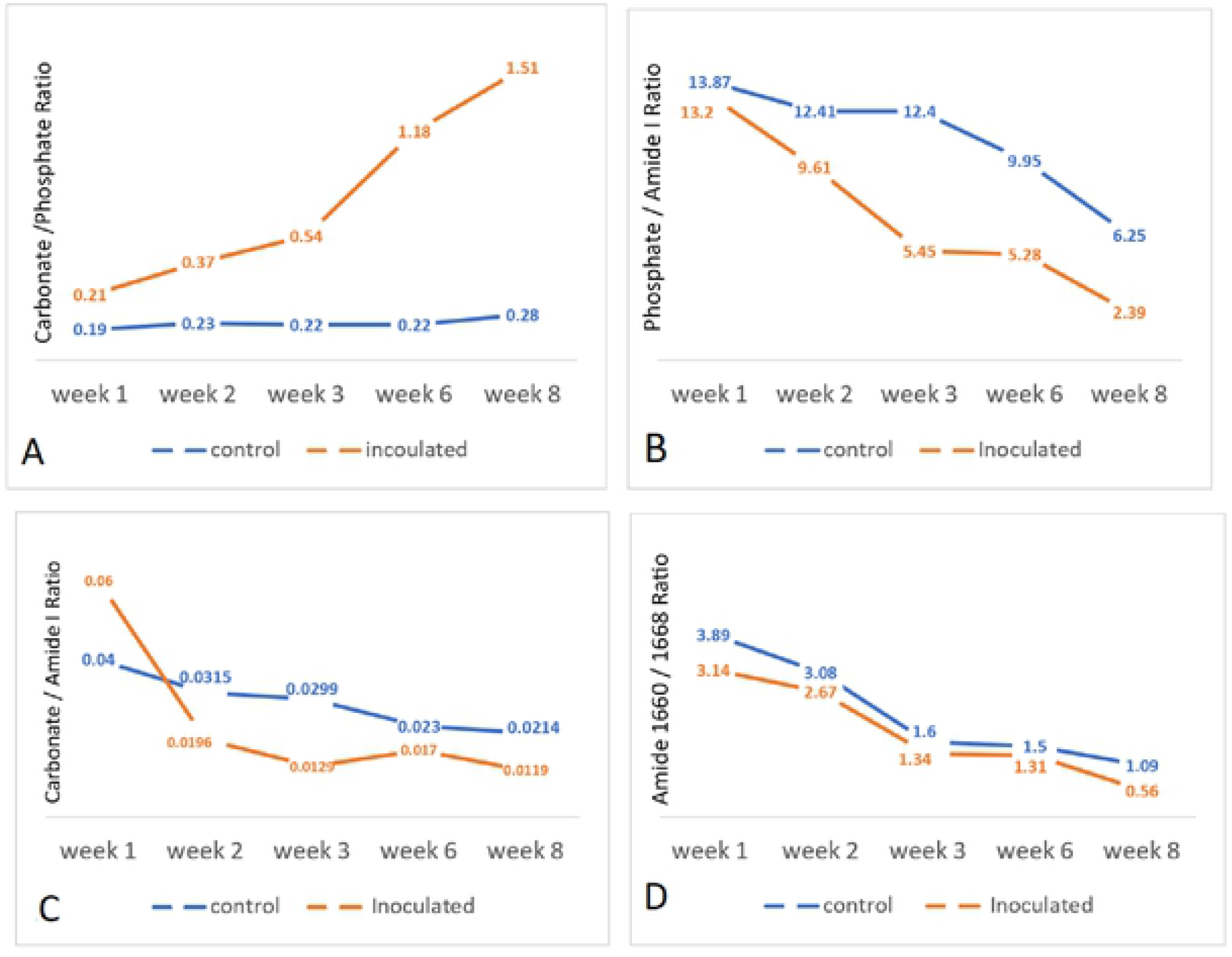
(A) Shows an increase in carbonate to phosphate ratio in inoculated samples compared to controls indicating changing bone crystallinity and transformation to a brittle matrix. (B&C) shows the reduction in mineral content relative to the organic material in inoculated samples compared to controls. (D) collagen crosslinking ratio shown as shift of the Raman bands at 1660cm–1 to 1668 cm–1 in inoculated samples compared to controls representing the destruction of collagen cross linkage

#### Scanning Electron Microscopy results

In weeks six and eight bone samples were examined using scanning electron microscopy (SEM) following appropriate preparation. SEM showed standard honeycomb trabecular appearance of cancellous bone in the control samples and destruction of the trabeculae in the inoculated specimens under different SEM magnifications.

## Discussion

The pathogenesis of osteomyelitis encompasses three agents: the pathogen, the host, and the local environment. Our focus in this experiment was to examine the ability of biofilm-forming pathogens to alter the bone structure in the absence of host defences. The first promulgation of biofilm hypothesis was in 1978. The theory stated that bacteria might grow predominantly in all nutrient sufficient environments in matrix-enclosed surface-associated groups which can protect it from host defences or antimicrobial agents^(9)^. It proposes that biofilm pathogens have the virulent activity where it can cause direct organic tissue damage.

However, the extent of direct bone resorption caused by the biofilm and its clinical significance remain unclear. Adam, J. et al. proposed mechanistic possibilities for the immediate bone destruction^(3)^. They suggested that virulence effect of planktonic bacteria start once it adheres to the bone surface. Then it starts to form the biofilm producing an extracellular polymeric slime matrix which can recruit and increase the adhesion of other bacteria. Later it forms mature biofilms which can cause bone cavitation and eventually bone fragments detached from the infected piece^(5)^. In chronic osteomyelitis, an avascular bone segment can harbour pathogenic biofilm which causes bone lysis and cavitation^(9)^. For example, the lysis and cavitation around prosthetic implants in chronic infection may help in the diagnosis when clinical symptoms are unclear ^(10, 11)^. This is similar to what we found in this study where the examined bones showed cavitation and destruction of the regular bony matrix which was evident under scanning electron microscopy. The findings of this experimental study indicated the ability of the pathogen biofilm to induce direct bone resorption similar to those seen in the clinical setting of osteomyelitis. It also showed the ability of Raman spectroscopy to pick up these changes. All tested inoculated bone samples were showing significant quantitative and qualitative structural damage induced by the *Staphylococcus aureus* biofilm compared to the control samples. Noticeable differences were found at later time points between the two groups in the ratio of phosphate to amide I ratio, carbonate to amide I ratio and amide 1660 to1668 ratio but was not statistically significant. We hypothesise that hydration can have effect on the bone quality, yet bacteria biofilm had significant bone destruction which was observed in the earlier weeks. However, at later weeks bacterial counts may have started dropping due to lack of nutrients in the media and therefore osteolytic activity was reduced.

However, the literature lacks sufficient details of the biofilm pathogens virulent factors and mechanisms which cause the direct tissue damage^(12)^. Therefore, comprehensive analysis of these negative factors might give a better understanding of how bone loss occurs and help in disease management.

## Conclusion

This experiment expands our understanding of bone destruction process during osteomyelitis. It also shows the direct effect of bacteria on bone in addition to the recognised destruction caused by osteoclastic and host cells.

## Limitation

The used Raman system (I-Raman Ex spectrometer, BWTek, Newark, USA) scans a spot size of 85um; therefore, an error of scanning of healthy spot in examined bone samples may have occurred, inadvertently. To minimise the effect of this potential error we used an average of multiple readings of the investigated samples. Also, the numbers were not large enough to show statistical significance of the changes that occur at molecular level.

## Notes

## Compliance with ethical standards

Conflict of interest: The authors declare that they have no conflict of interest.

Ethical approval: The approval of the study was obtained from Sultan Qaboos University medical ethical committee *(REF. NO. SQU-EC1039114).*

Informed consent: written informed consent was obtained from the patients

